# Genome-wide Association Study of Alcohol Consumption and Use Disorder in Multiple Populations (N = 274,424)

**DOI:** 10.1101/527929

**Authors:** Henry R. Kranzler, Hang Zhou, Rachel L. Kember, Rachel Vickers Smith, Amy C. Justice, Scott Damrauer, Philip S. Tsao, Derek Klarin, Daniel J. Rader, Regeneron Genetics Center Research Team, Zhongshan Cheng, Janet P. Tate, William C. Becker, John Concato, Ke Xu, Renato Polimanti, Hongyu Zhao, Joel Gelernter, on behalf of the VA Million Veteran Program

## Abstract

Although alcohol consumption level and alcohol use disorder (AUD) diagnosis are both moderately heritable, their genetic risks and overlap are not well understood. We conducted genome-wide association studies of these traits using longitudinal Alcohol Use Disorder Identification Test-Consumption (AUDIT-C) scores (reflecting alcohol consumption) and AUD diagnoses from electronic health records (EHRs) in a single, large multi-ancestry Million Veteran Program sample. Meta-analysis across population groups (N = 274,424) identified 18 genome-wide significant loci, 5 of which were associated with both traits and 13 with either AUDIT-C (N = 8) or AUD (N = 5). A significant genetic correlation between the traits reflects this overlap. However, downstream analyses revealed biologically meaningful points of divergence. Cell-type group partitioning heritability enrichment analyses indicated that central nervous system was the most significant cell type for AUDIT-C and the only significant cell type for AUD. Polygenic risk scores (PRS) for both traits were associated with alcohol-related disorders in two independent samples. Genetic correlations for 188 non-alcohol-related traits were significantly different for the two traits, as were the phenotypes associated with the traits’ polygenic risk scores. We conclude that EHR-derived, longitudinal, repeated measures of alcohol consumption level and AUD diagnosis can facilitate genetic discovery and help to elucidate the relationship between drinking level and AUD risk. Finally, although heavy drinking is a key risk factor for AUD, it is not a sufficient cause of the disorder.

---

Excessive alcohol consumption is associated with a host of adverse medical, psychiatric, and social consequences. Globally, in 2012, about 3.3 million or 5.9% of all deaths,139 million disability-adjusted life years, and 5.1% of the burden of disease and injury, were attributable to alcohol consumption, with the magnitude of harm determined by the volume of alcohol consumed and the drinking pattern^1^. Regular heavy drinking is the major risk factor for the development of an alcohol use disorder (AUD), a chronic, relapsing condition characterized by impaired control over drinking^2^. Independent of AUD, heavy drinking has a multitude of adverse medical consequences. Identifying factors that contribute to drinking level and AUD risk could advance efforts to prevent, identify, and treat both medical and psychiatric problems related to alcohol.

Many different alcohol-related phenotypes have been used to investigate genetic risk, including formal diagnoses, such as alcohol dependence [e.g., based on the Diagnostic and Statistical Manual of Mental Disorders, 4^th^ edition (DSM-IV)^3^] and screening tests that measure alcohol consumption and alcohol-related problems (e.g., the Alcohol Use Disorders Identification Test (AUDIT). The AUDIT, a 10-item, self-reported test developed by the World Health Organization as a screen for hazardous and harmful drinking^4,5^ has been used for genome-wide association studies (GWASs) both as a total score^6–8^, and as the AUDIT-Consumption (AUDIT-C) and AUDIT-Problems (AUDIT-P) sub-scores^8^. The three-item AUDIT-C measures the frequency and quantity of usual drinking and the frequency of binge drinking, while the 7-item AUDIT-P measures alcohol-related problems.

Twin and adoption studies have shown that half of the risk of alcohol dependence, a subtype of AUD, is heritable^9^. The SNP heritability of alcohol dependence in a family-based, European-American (EA) sample was 16%^10^ and 22% in an unrelated African-American (AA) sample^11^. In the meta-analysis of data from the UK Biobank (UKBB) and 23andMe, the SNP heritability of the total AUDIT was estimated to be 12%, while for the AUDIT-C and AUDIT-P it was 11% and 9%, respectively)^8^.

In 12 genome-wide association studies (GWASs) of alcohol dependence (most of which used a binary DSM-IV diagnosis^3^) published between 2009 and 2014^12^, the only consistent genome-wide significant (GWS) findings were for single nucleotide polymorphisms (SNPs) in genes encoding the alcohol metabolizing enzymes. Similarly, in a recent meta-analysis of 14,904 individuals with alcohol dependence and 37,944 controls, which was stratified by genetic ancestry (European, N = 46,568; African; N = 6,280), the only GWS findings were two independent *ADH1B* variants. In addition, there were significant genetic correlations seen with 17 phenotypes, including psychiatric (e.g., schizophrenia, depression), substance use (e.g., smoking and cannabis use), social (e.g., socio-economic deprivation) and behavioral (e.g., educational attainment) traits^13^.

Alcohol metabolizing enzyme genes have also been associated with mean or maximal alcohol consumption levels, potential intermediate phenotypes for alcohol dependence^14–19^. In a meta-analysis of GWASs (N > 105,000 European subjects), *KLB* was associated with alcohol consumption^20^. A GWAS of alcohol consumption in the UK Biobank sample^21^ identified GWS associations at 14 loci (8 independent), including three alcohol metabolizing genes on chromosome 4 (*ADH1B, ADH1C*, and *ADH5*), an intergenic SNP on chromosome 4, and *KLB*, replicating the prior meta-analytic findings. Novel risk genes identified in this study included *GCKR, CADM2*, and *FAM69C*.

A GWAS of the AUDIT in nearly 8,000 individuals failed to identify any GWS loci^6^. A GWAS of the AUDIT from 23andMe in 20,328 European ancestry participants also failed to yield GWS results^7^, although meta-analysis of the AUDIT in the UKBB and 23andMe samples identified 10 associated risk loci, including novel associations to *JCAD* and *SLC39A13*^8^. In addition to the total AUDIT-C score, the meta-analysis included GWASs for the AUDIT-C and AUDIT-P, which showed significantly different patterns of association across a number of traits, including psychiatric disorders. Specifically, genetic correlations between schizophrenia, major depressive disorder, and obesity (among others) were negative for AUDIT-C and positive for AUDIT-P.

In the present study, we evaluated the independent and overlapping genetic contributions to AUDIT-C and AUD in a single large multi-ancestry sample from the Million Veteran Program (MVP)^22^. Large-scale biobanks such as the MVP offer the potential to link genes to health-related traits documented in the electronic health record (EHR) with greater statistical power than can ordinarily be achieved in prospective studies^23^. Such discoveries improve our understanding of the etiology and pathophysiology of complex diseases and their prevention and treatment. To that end, we used a common data source—longitudinal repeated measures of alcohol-related traits from the national Veterans Health Administration (VHA) EHR—to obtain the mean, age-adjusted AUDIT-C score and International Classification of Diseases (ICD) alcohol-related diagnosis codes over more than 11 years of care^24^. We then conducted a GWAS of each trait followed by downstream analysis of the findings in which we constructed polygenic risk scores (PRS) for both traits and showed that they were associated with alcohol-related disorders in two independent samples. The availability of data on alcohol consumption from the AUDIT-C and a formal diagnosis of AUD from the EHR enabled us to examine the relationship between these key alcohol-related traits in a single, well-phenotyped sample and to compare the findings for these traits more systematically than has previously been possible.

## Results

### Principal Components Analysis (PCA)

We differentiated participants genetically into 5 populations (see Methods, Supplementary Figure 1) and removed outliers. There was a high degree of concordance (Supplementary Figure 2) between the genetically defined populations and the self-reported groups for European Americans (EAs, 95.6% were self-reported Non-Hispanic white) and African Americans (AAs, 94.5% were self-reported Non-Hispanic black). Concordance ranged from 53.1-81.6% in the other 3 population groups.

### GWAS Analyses

#### AUDIT-C

The GWAS for AUDIT-C (Figure 1a, Table 1 and Supplementary Tables 1 and 2) identified 13 independent loci in EAs, 2 in AAs, 1 in LAs (Hispanic and Latino Americans) and 1 in EAAs (East Asian Americans) (Supplementary Figures 4 and 5). Meta-analysis across the 5 populations (see Methods) also yielded 13 independent loci, 5 of which were previously associated with a self-reported measure of alcohol consumption: *GCKR*^21^, *KLB*^20,21^, *ADH1B*^18,21^, *ADH1C*^21^, and *SLC39A8*^8^. The 8 novel trans-population signals for AUDIT-C included *VRK2* (Vaccinia related kinase 2), *DCLK2* (Doublecortin like kinase 2), *ISL1* (ISL LIM Homeobox 1), *FTO* (Alpha-Ketoglutarate Dependent Dioxygenase), *IGF2BP1* (Insulin like growth factor 2 MRNA binding protein 1), *PPR1R3B* (Protein phosphatase 1 regulatory subunit 3B), *BRAP* (BRCA1 associated protein), *BAHCC1* (BAH domain and coiled-coil containing 1), and *RBX1* (Ring-box 1). *BAHCC1* and *RBX1* were GWS only in the trans-population meta-analysis, the results of which were driven largely by the findings in EAs, who comprised 73.5% of the total MVP sample.

**Figure 1.**
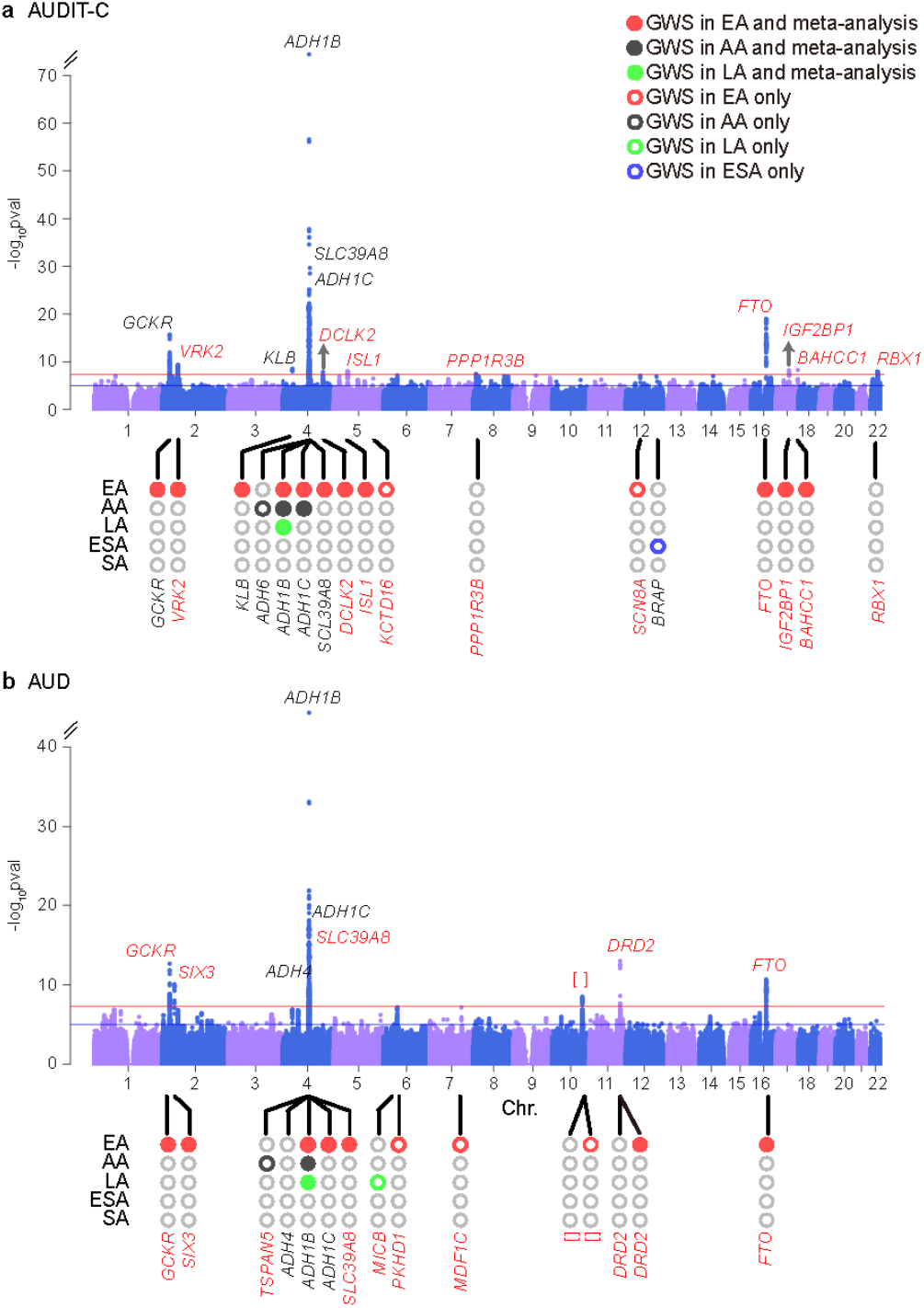
Manhattan plots for age-adjusted mean AUDIT-C score and AUD diagnosis. a) Manhattan plot of the genome-wide association meta-analysis of AUDIT-C across all 5 populations (N = 278,424). b) Manhattan plot of the genome-wide association meta-analysis of AUD across 5 populations (55,985 cases and 224,014 controls). Red lines show the genome-wide significance level (5.0 × 10^−8^). EA: European American; AA: African American; LA: Hispanic or Latino; EAA: East Asian American; SAA: South Asian American. Labeled genes at the top of the peaks indicate completely independent signals after conditional analysis in meta-analysis. Population-specific loci are labeled at the bottom of the circles in the lower part of each figure. Genes labeled in red have not previously been reported for the trait. []: no genes within 500kb to the lead SNP.

**Table 1.**
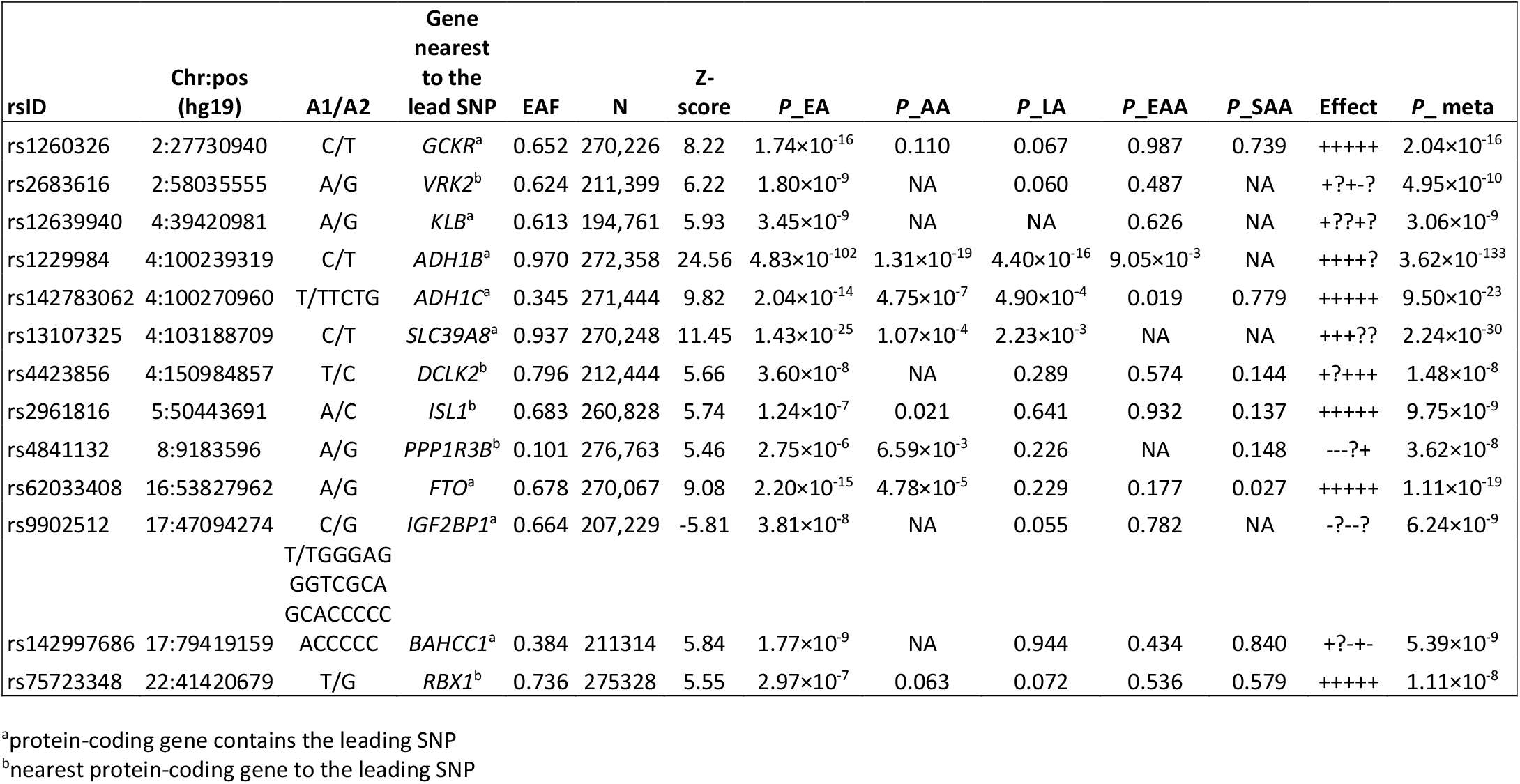
Genome-wide significant associations for AUDIT-C in the trans-population meta-analysis. The loci shown represent completely independent signals after conditioning analyses. A1, effect allele; A2, other allele; EAF, effective allele frequency; EA, European American; AA, African American; LA, Latino American; EAA, East Asian American; SAA, South Asian American.

#### AUD

The GWAS for AUD (Figure 1b, Table 2 and Supplementary Tables 1 and 3) identified 10 independent loci in EAs, 2 in AAs, and 2 in LAs (Supplementary Figures 6 and 7). Meta-analysis across the 5 populations yielded 10 independent loci, including 3 previously associated with alcohol dependence^25^—*ADH1B, ADH1C*, and *ADH4*—and 7 loci not previously associated with an AUD diagnosis: *GCKR, SIX3* (SIX Homeobox 3), *SLC39A8, DRD2* (Dopamine Receptor D2: rs4936277 and rs61902812, which were independent), chr10q25.1 (rs7906104), and *FTO*. Five loci were significant in both the AUDIT-C and AUD GWASs (Supplementary Figure 8): *ADH1B, ADH1C, FTO, GCKR*, and *SLC39A8*. The trans-population GWS findings for AUD were also driven largely by the findings in EAs.

**Table 2.**
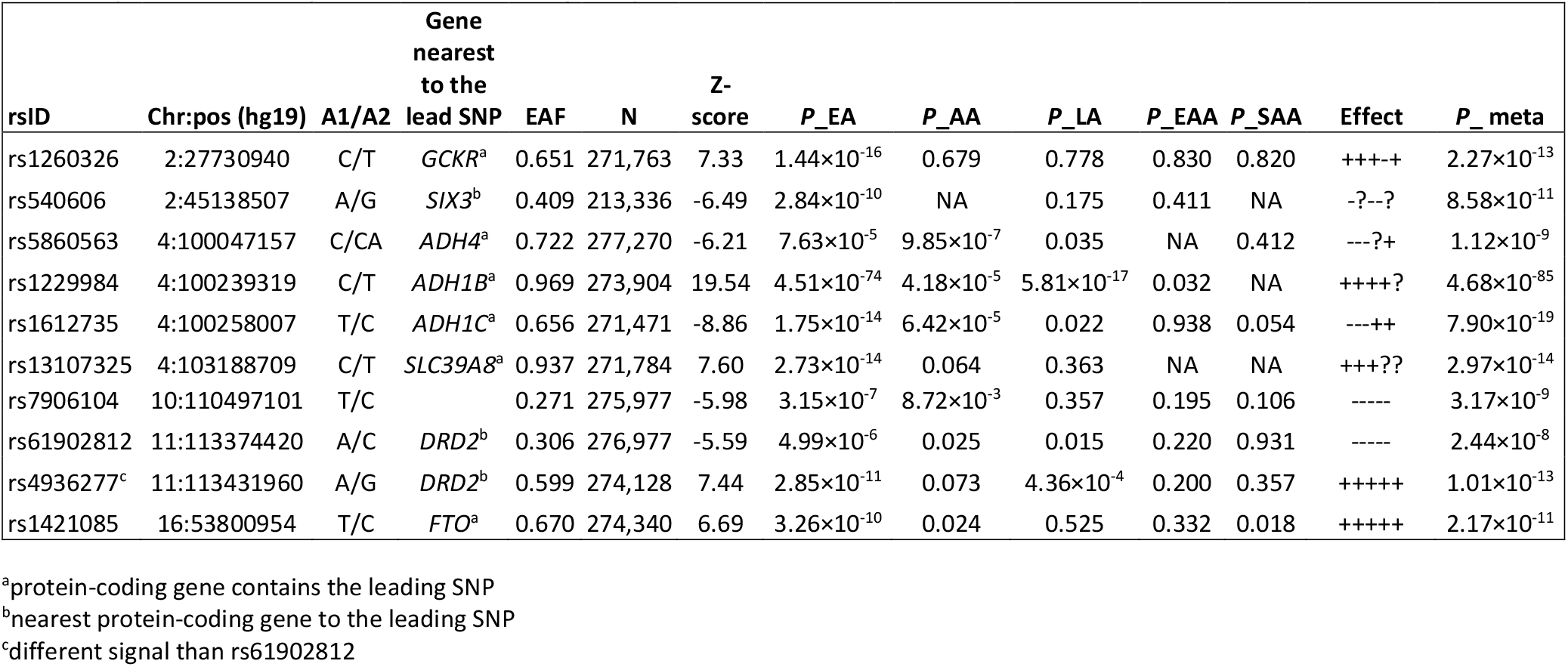
Genome-wide significant associations for AUD in the trans-population meta-analysis. The loci shown represent completely independent signals after conditioning analyses.

#### Sex-specific Effects

The GWAS findings largely reflect male-specific signals due to the predominantly male sample (Supplementary Tables 1). However, sex-stratified GWAS also identified two female-specific signals for AUDIT-C (Supplementary Table 4, Supplementary Figures 9 and 10) and one for AUD (Supplementary Table 5, Supplementary Figures 11 and 12).

#### Conditional Analyses

For AUDIT-C, when associations for the 7 LD-pruned GWS SNPs on chromosome 4q23-q24 in EAs were conditioned on rs1229984, the most significant functional SNP in the region in that population, the only independent signal (using a Bonferroni-corrected p-value < 0.05) was for rs1229978, near *ADH1C* (Supplementary Table 6). For AUD, when associations for the 4 LD-pruned GWS SNPs in EAs were conditioned on rs1229984, the only independent signal in that region was for rs1154433 near *ADH1C*. In the trans-population meta-analysis, rs5860563 was independent when conditioned on rs1229984 in EAs and on rs2066702 in AAs, the most significant functional SNP in the region in AAs (Supplementary Table 7).

To elucidate further the genetic differences between AUDIT-C and AUD, we conducted a GWAS of each phenotype with the other phenotype as a covariate. A GWAS of AUDIT-C with AUD as a covariate identified 10 GWS loci in EAs and 2 GWS loci in AAs (Supplementary Table 8). In both EAs and AAs, all loci overlapped with the GWS findings for AUDIT-C alone. A GWAS of AUD that included AUDIT-C as a covariate identified 5 GWS loci in EAs and one in AAs (Supplementary Table 9). Among EAs, 4 of the loci were the same as for AUD, the only non-overlapping finding being *DIO1* (Iodothyronine Deiodinase 1). In AAs, *ADH1B* remained significant for AUD when accounting for AUDIT-C, but *TSPAN5* did not.

Using a sign test, we found that most SNPs had the same direction of effect for AUDIT-C and AUD, consistent with the high genetic correlation between the traits. For SNPs with p-value < 1×10^−6^ the sign concordance between the two traits was 98.7% in EAs and 100% in the other four, smaller population groups.

#### BMI-Adjusted GWAS

Because *FTO* was GWS for both AUDIT-C and AUD, we repeated the two GWASs correcting for BMI. Among the top SNPs associated with AUDIT-C and AUD, most remained GWS after correction for BMI, though the significance level of some changed (Supplementary Tables 10 and 11). *FTO* SNPs became only nominally significant for both alcohol-related traits: the p-value for the lead SNP for AUDIT-C, rs9937709, decreased in significance from 5.53 × 10^−14^ to 1.42 × 10^−5^ and for the lead SNP for AUD, rs11075992, it decreased in significance from 3.22 × 10^−10^ to 3.02 × 10^−5^. In contrast, with correction for BMI, some signals increased, e.g., for rs1260326 in *GCKR* the p-value increased in significance from p = 2.04 × 10^−16^ to p = 2.91 × 10^−19^ for AUDIT-C and from p = 2.27 × 10^−13^ to p = 1.71 × 10^−14^ for AUD. Similarly, rs1229984 in *ADH1B* increased in significance from p = 3.62 × 10^−133^ to p = 9.81 × 10^−145^ for AUDIT-C and from p = 4.68 × 10^−85^ to p = 3.85 × 10^−89^ for AUD.

### Gene-based Analyses

#### AUDIT-C

For AUDIT-C score, gene-based association analyses identified 31 genes in EAs that were GWS (p < 2.69 × 10^−6^), 3 in AAs, 1 in LAs, and 2 in EAAs (Supplementary Figure 13), including many of the loci in the SNP-based analyses for that trait. The unique genes in EAs included *C4orf17, ZNF512, MTTP, TBCK*, and *MCC*. For AUDIT-C, the loci that were not GWS in the SNP-based analyses included *EIF4E* in AAs, *MAP2* in LAs, and *LOX* and *MYL2* in EAAs.

#### AUD

For AUD, we identified 23 GWS genes in EAs, 5 in AAs and 1 in LAs (Supplementary Figure 14), many of which were GWS loci in the SNP-based analyses for that trait. For AUD, the loci in EAs that were not GWS in the SNP-based analyses were *KRTCAP3, TRMT10A, ZNF512, DCLK2, MTTP*, and *MCC*. In AAs, *EIF4E, ADH4*, and *METAP1* were GWS for AUD, while *ADGRB2* was the only GWS locus in LAs.

#### Pathway and Biological Enrichment Analyses

Using Functional Mapping and Annotation (FUMA)^26^ software to investigate the pathway or biological process enrichment with summary statistics as input and FDR correction for multiple testing, we found multiple reactome and Kyoto Encyclopedia of Genes and Genomes (KEGG) pathways that were significantly enriched for AUDIT-C (Supplementary Table 12, Supplementary Figure 15) and AUD (Supplementary Table 13, Supplementary Figure 16) in each population. The most significant pathway was reactome ethanol oxidation for both traits in both EAs and AAs. Multiple GO biological processes were enriched for AUDIT-C (Supplementary Table 14, Supplementary Figure 17) and AUD (Supplementary Table 15, Supplementary Figure 18), including ethanol and alcohol metabolism. Enrichments for chemical and genetic perturbation gene sets and for the GWAS catalog for both traits are shown in Supplementary Tables 16-19 and Supplementary Figures 19-22.

#### Heritability Estimates

We used linkage disequilibrium score regression (LDSC)^27^ (see Methods) to estimate SNP-based heritability 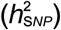 in EAs and AAs, where sample sizes were large enough to provide robust estimates for each trait (Figure 2a). For AUDIT-C, the 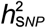 was 0.068 (se = 0.005) in EAs: 0.068 (se = 0.005) in males and 0.099 (se = 0.037) in females. In AAs, the 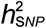 was 0.062 (se = 0.016): 0.058 (se = 0.018) in males. For AUD, the 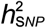 was 0.056 (se = 0.004) in EAs: 0.054 (se = 0.004) in males and 0.110 (se = 0.038) in females. The 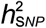 for AUD was 0.100 (se = 0.022) in AAs; 0.104 (se = 0.023) in males. Robust estimates of 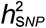 were unavailable in AA females due to the small sample size.

**Figure 2.**
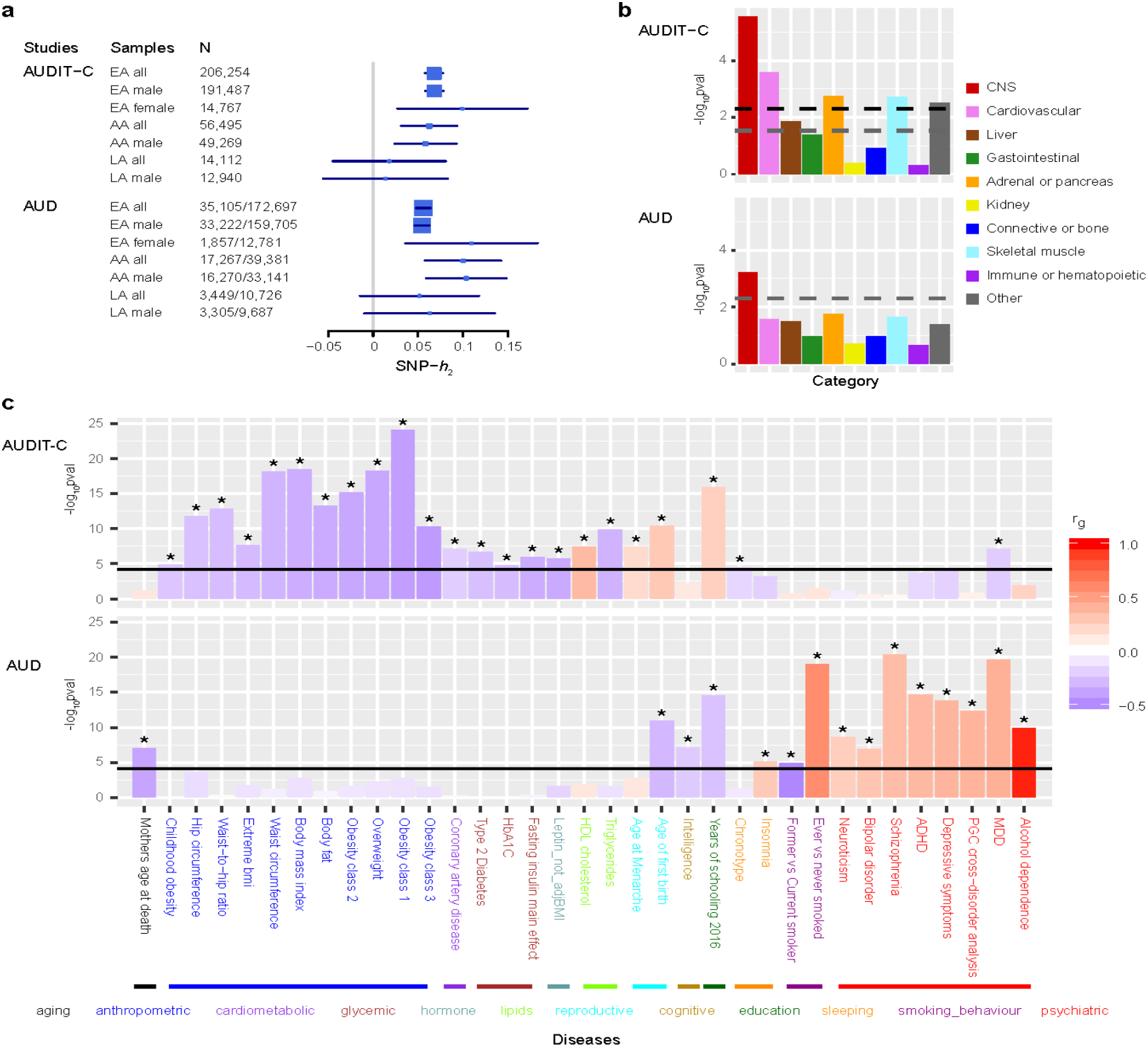
Heritability estimate, partitioning enrichments of heritability, and genetic correlation analyses for age-adjusted mean AUDIT-C score and AUD diagnosis using LD score regression. a) SNP-based heritability for AUDIT-C and AUD in the three populations and sex-stratified samples adequate in size for the analysis. b) Partitioned heritability enrichment of cell type groups for AUDIT-C and AUD. Ten cell types tested were corrected for multiple testing. The black dashed line is the cutoff for Bonferroni-corrected significance. The gray dashed lines are the cutoff for FDR < 0.05. c) Genetic correlations with other traits. Data from 714 publicly available datasets (221 published and 493 unpublished from UK Biobank) were tested and corrected for multiple comparisons. The traits presented are for published data. Black lines are the cutoff for Bonferroni-corrected significance, with asterisks showing traits significant after correction. The traits are grouped into different categories and sorted by the genetic correlations with AUDIT-C (upper panel) or AUD (lower panel). CNS: central nervous system. ADHD: attention deficit hyperactivity disorder. MDD: major depressive disorder.

In the analysis of stratified heritability enrichment using LDSC^28^ (see Methods), several cell line functional enrichments were significant (false discovery rate (FDR) < 0.05) for AUDIT-C (Supplementary Table 20) and AUD (Supplementary Table 21). Cell-type group partitioning heritability enrichment analyses indicated that central nervous system (CNS) was the most significant cell type for AUDIT-C (Figure 2b, upper panel; Supplementary Table 22) and the only significant cell type for AUD (Figure 2b, bottom panel; Supplementary Table 23). Enrichments for AUDIT-C were also detected for cardiovascular, adrenal or pancreatic, skeletal muscle, other, and liver cell types in descending order of significance. We also tested the heritability enrichments using data from gene expression and chromatin to identity disease-related tissues or cell types^29^ (Supplementary Tables 24-33). We found a few epigenetic features in brain tissues—e.g., H3K4me1, H3K4me3, and DNase–that were significantly enriched for each trait.

#### Genetic Correlations

We estimated the genetic correlation (*r*_g_) between different datasets or populations using LDSC^30^. The *r*_g_ between AUDIT-C and AUD was 0.522 (se = 0.038, p = 2.40 × 10^−42^) in EAs and 0.930 (se = 0.122, p = 1.85 × 10^−14^) in AAs (Supplementary Table 34). The *r*_g_ between EA males and EA females was 0.815 (se = 0.156, p = 1.69 × 10^−7^) for AUDIT-C, and 0.833 (se = 0.142, p = 4.16 × 10^−9^) for AUD.

After Bonferroni correction, 179 traits or diseases were genetically correlated with AUDIT-C (Figure 2c, upper panel; Supplementary Table 35). AUDIT-C was positively genetically correlated with lipids (e.g., HDL cholesterol concentration: *r*_g_ = 0.361, p = 3.39 × 10^−8^), reproductive traits (e.g., age at menarche: 0.190, p = 4.20 × 10^−8^), and years of education (*r*_g_ = 0.248, p = 1.40 × 10^−16^) and negatively correlated with anthropometric (e.g., body mass index (BMI): *r*_g_ = −0.350, p = 3.25 × 10^−19^), cardiometabolic (e.g., coronary artery disease: *r*_g_ = −0.212, p = 8.28 × 10^−8^), glycemic (e.g., Type 2 diabetes: *r*_g_ = −0.273, p = 2.34 × 10^−7^), lipids (e.g., triglyceride concentration: *r_g_* = −0.325, p = 1.29 × 10^−10^), and psychiatric (e.g., major depressive disorder (MDD) (*r*_g_ = −0.216, p = 7.72 × 10^−8^) traits. After correction, 111 traits or diseases were genetically associated with AUD (Figure 2c bottom panel; Supplementary Table 36), including positive genetic correlations with sleep disturbance (e.g., insomnia: *r*_g_ = 0.280, p = 7.43 × 10^−6^), ever having smoked (*r*_g_ = 0.581, p = 9.19 × 10^−20^), and multiple psychiatric disorders (e.g., alcohol dependence: *r_g_* = 0.965, p = 1.21 × 10^−10^, MDD: *r*_g_ = 0.406, p = 2.19 × 10^−20^) and negative genetic correlations with aging-related factors (e.g., mother’s age at death: *r*_g_ = −0.390, p = 8.09 × 10^−8^), intelligence (*r*_g_ = −0.226, p = 6.79 × 10^−8^), years of education (*r*_g_ = −0.263, p = 2.88 × 10^−15^), and quitting smoking (*r*_g_ = −0.517, p = 1.12 × 10^−5^).

We tested the difference between genetic correlations for AUDIT-C and AUD using a two-tailed z-test. After correction for 714 tested traits, the genetic correlations for 188 traits showed significant differences between the two alcohol-related traits (Supplementary Table 37).

We explored trait and disease associations for AUDIT-C-adjusted for AUD, and AUD-adjusted for AUDIT-C, and found that the genetic correlations between the alcohol-related traits and other phenotypes did not differ substantially from the unadjusted ones (Supplementary Tables 38 and 39). Additionally, we explored genetic correlations between AUDIT-C-adjusted for BMI (Supplementary Table 40), and AUD-adjusted for BMI (Supplementary Table 41). Most of the genetic correlations for AUDIT-C-adjusted for BMI did not differ substantially from the unadjusted ones, except for anthropometric traits, where the negative correlation was attenuated (although still significant). Significant genetic correlations for AUD-adjusted for BMI did not differ substantially from those for AUD alone. We also explored prior GWAS associations for the GWS SNPs from AUDIT-C and AUD analyses and found associations with other phenotypes for 5 of them (Supplementary Table 42).

#### Polygenic Risk Scores

We examined PRS generated from the AUDIT-C and AUD GWASs in three samples (Supplementary Figures 23-26). First, in a “hold-out” MVP sample of EAs and AAs (described in Methods), AUDIT-C and AUD PRS were significantly associated with both AUDIT-C and AUD phenotypes (Supplementary Tables 43 and 44). Lower p-value thresholds of AUDIT-C PRS were associated with AUDIT-C score and AUD diagnosis codes, with the most significant being 1 × 10^−7^ (EA AUDIT-C: β = 0.088, p = 1.43 × 10^−44^; EA AUD: β = 0.137, p =3.03 ×10^−30^; AA AUDIT-C: β = 0.094, p =2.82 × 10^−17^; AA AUD: β = 0.110, p = 1.3 × 10^10^). All p-value thresholds for AUD PRS were associated with both AUDIT-C score and AUD diagnosis codes, with the most significant being 1 × 10^−7^ for EAs (AUDIT-C: β = 0.095, p = 8.98 × 10^−51^; AUD: β = 0.147, p = 6.02 × 10^−34^) and 1 × 10^−7^ for AAs (AUDIT-C: β = 0.066, p = 3.69 × 10^−9^; AUD: β = 0.098, p = 1.09 × 10^−9^).

Second, in an independent sample from the Penn Medicine BioBank, AUDIT-C and AUD PRS were significantly associated with alcohol-related disorders and alcoholism phecodes (see Methods and Supplementary Tables 45 and 46). In EAs, higher AUDIT-C risk scores significantly increased the likelihood of alcohol-related disorders and alcoholism at multiple p-value thresholds, with the most significant being 1 × 10^−7^ (β = 0.278, p = 0.0013) and 1 × 10^−6^ (β = 0.245, p = 0.0074), respectively. In AAs, at a p-value threshold of 1 × 10^−7^, AUDIT-C risk scores were non-significantly associated with risk of alcohol-related disorders (β = 0.210, p = 0.064) but significantly associated with alcoholism (β = 0.400, p = 0.0051). In both populations, AUD risk scores were significantly associated with both alcohol-related disorders and alcoholism. In EAs, the most significant p-value threshold was 1 × 10^−4^ (alcohol-related disorders: β = 0.254, p = 0.0006; alcoholism: β = 0.229, p = 0.0062), while in AAs, the most significant p-value threshold was 1 × 10^−6^ (alcohol-related disorders: β = 0.306, p = 0.006; alcoholism: β = 0.440, p = 0.0007).

Third, in the Yale-Penn study sample^25^, an independent sample ascertained for substance use disorders, the PRS of AUDIT-C and AUD were significantly associated with DSM-IV alcohol dependence criterion counts (see Methods and Supplementary Tables 47 and 48). In EAs, all AUDIT-C risk scores were significantly associated with the criterion count, with the most significant p-value threshold being 1 × 10^−7^ (β = 1.029, p = 6.67 × 10^−13^). Similarly, all AUD risk scores were significantly associated with the criterion count, the most significant p-value threshold being 1 × 10^−6^ (β = 1.144, p = 1.86 × 10^−16^). In AAs, all but one AUDIT-C risk score and all AUD risk scores were significantly associated with the alcohol dependence criterion count, the most significant p-value threshold being 1 × 10^−7^ (AUDIT-C: β = 0.829, p = 1.18 × 10^−11^; AUD: β = 0.502, p = 4.62 × 10^−8^).

#### Secondary phenotypic associations

To identify secondary phenotypes associated with AUDIT-C or AUD, we performed a phenome-wide association analysis (PheWAS) of the AUDIT-C and AUD PRS (p-value threshold = 1 × 10^−7^ and all SNPs) in the MVP “hold-out” sample (Supplementary Tables 49 and 50). In EAs, the AUDIT-C PRS was significantly associated with an increased risk of alcoholic liver damage, and nominally associated with a decreased risk of hyperglyceridemia. No significant associations were found for AAs. The AUD PRS was significantly associated with an increased risk of tobacco use disorder in both EAs and AAs, and in EAs with multiple psychiatric disorders, including major depression, bipolar disorder, anxiety, and schizophrenia.

## Discussion

We report here the largest, most diverse single-sample, alcohol-related GWAS to date, including 274,424 MVP participants from five population groups—European American, African American, Latino American, East Asian American, and South Asian American—using two EHR-derived phenotypes: age-adjusted AUDIT-C score and AUD diagnostic codes. In addition to the large number of EAs, the sample included the largest numbers, by far, of African-American and Latino-American participants in a GWAS of alcohol-related traits. Trans-population meta-analyses identified 13 independent GWS loci for AUDIT-C, 8 of which are novel, and 10 independent GWS loci for AUD, including 7 novel findings. For AUDIT-C, in addition to the loci identified in the SNP-based analyses, there were 31 GWS genes in EAs, 3 in AAs, 1 in LAs, and 2 in EAAs. For AUD, in addition to the loci identified in the SNP analyses, there were 23 GWS genes identified in EAs, 5 in AAs, and 1 in LAs.

Using both AUDIT-C scores and AUD diagnoses enabled us to examine the relations between these key alcohol-related traits. The findings underscored the utility of using an intermediate trait, such as alcohol consumption, for genetic discovery. Five of the 13 loci associated with AUDIT-C score, a measure of alcohol consumption, including the two most commonly identified alcohol metabolism genes (*ADH1B* and *ADH1C*) and three highly pleiotropic genes (*GCKR, SLC39A8*, and *FTO*), contributed to AUD risk. Of the 10 loci that were GWS for AUD, half also were associated with AUDIT-C score, while half were uniquely associated with the AUD diagnosis: *ADH4, SIX3*, a variant on chr10q25.1 and 2 variants in *DRD2*.

In addition to multiple overlapping variants for AUDIT-C and AUD, we found a moderate-to-high genetic correlation between the traits: 0.522 in EAs and 0.930 in AAs. This population difference may reflect a bias in the assignment of AUD diagnoses by clinicians (e.g., in the context of a high AUDIT-C score, clinicians could be less likely to assign an AUD diagnosis to EAs than AAs). Another factor relevant to this difference is the smaller number of AAs, which despite a higher *r*_g_, yielded a larger standard error. The genetic similarity between these alcohol-related traits is consistent with twin studies of alcohol dependence and alcohol consumption^31,32^. These findings are also consistent with the PRS analyses in the MVP sample, where both AUDIT-C and AUD PRS were associated with AUDIT-C and AUD phenotypes. Both traits also predicted multiple alcohol-related phenotypes in independent datasets, including alcohol dependence criteria in the Yale-Penn sample. However, there was a smaller effect of AUDIT-C PRS scores than AUD PRS scores on alcohol-related disorders and alcohol dependence. This is in line with findings from the meta-analysis of UKBB and 23andMe data, where the genetic correlation with alcohol dependence was nominally greater for AUDIT-P scores (*r*_g_ = 0.63) than AUDIT-C scores (*r*_g_ = 0.33)^8^.

Despite the significant genetic overlap between the AUDIT-C and AUD, downstream analyses revealed biologically meaningful points of divergence. The AUDIT-C yielded many GWS findings that did not overlap with those for AUD, which reflects genetic independence of the traits. This broadens our previous observations using SNPs in *ADH1B*, in which we validated the AUDIT-C score as an alcohol-related phenotype^33^. In that study, after accounting for the effects of AUDIT-C score, AUD diagnoses accounted for unique variance in the frequency of the minor alleles.

Evidence of genetic independence between the two traits was most striking in the differences between the genetic correlation analyses. After correction, genetic correlations for 188 traits differed significantly (some in opposite directions) between AUDIT-C and AUD. Notably, these included a negative association of AUDIT-C with anthropometric traits, including BMI; coronary artery disease; and glycemic traits, including Type 2 diabetes. The negative genetic correlation with coronary artery disease is consistent with some epidemiological findings that alcohol consumption protects against some forms of cardiovascular disease^34^. AUDIT-C was positively genetically correlated with overall health rating, HDL cholesterol concentration, and years of education, findings that are consistent with prior literature showing genetic correlation of these traits with alcohol consumption^7,8,21^. AUD was significantly genetically correlated with 111 traits or diseases, including negative genetic correlations with intelligence, years of education and quitting smoking, and positive genetic correlations with insomnia, ever having smoked and most psychiatric disorders, findings that are consistent with phenotypic associations in the epidemiological literature^35–37^ and genetic correlations reported from the UKBB and 23andMe GWASs and their meta-analysis^7,8,21^. Further, in the MVP sample, the AUD PRS was significantly positively associated with tobacco use and multiple psychiatric disorders, whereas the AUDIT-C PRS was not. Taken together, these findings suggest that AUD and alcohol consumption, measured by AUDIT-C, are related but distinct phenotypes, with AUD being more closely related to other psychiatric disorders, and AUDIT-C with some positive health outcomes.

Although the protective effects of moderate drinking are controversial, we found that alcohol consumption in the absence of genetic risk for AUD may protect from cardiovascular disease, diabetes mellitus, and major depressive disorder. In contrast, individuals with genetic risk for AUD are at elevated risk for some adverse secondary phenotypes, including insomnia, smoking, and other psychiatric disorders. However, because individuals who have had health problems resulting from drinking are more likely to reduce or stop drinking by middle age or under-report their alcohol consumption, an alternative explanation for the opposite genetic associations is the “sick quitter” phenomenon^38^, which might be operating in an older clinical sample in which a large proportion report current abstinence (reflected in an AUDIT-C score of 0). For this complex set of genetic associations to be useful in informing clinical recommendations on safe levels of alcohol consumption, it will be necessary to elucidate the mechanisms underlying these findings.

Both phenotypes showed cell type-specific enrichments for CNS. Other relevant cell types for AUDIT-C, but not for AUD, included cardiovascular, adrenal or pancreas, liver, and musculoskeletal. Thus, although heavy drinking is prerequisite to the development of AUD, the latter is a polygenic disorder and variation in genes expressed in the CNS (e.g., *DRD2*) may be necessary for individuals who drink heavily to develop AUD. As a binary trait, AUD provided less statistical power to identify genetic variation than the ordinal AUDIT-C score, but the multiple GWS findings unique to AUD argue against that as an explanation for the non-overlapping GWS findings for the two traits.

The VHA EHR provided a rich source of phenotypic data. These included mean age-adjusted AUDIT-C scores, which are more stable than measures at a single point in time (more likely reflecting “traits” rather than “states”) and contrast with meta-analytic studies that may use phenotypes reflecting the lowest-common denominator among the studies comprising the sample. However, our analyses were limited by our reliance on the AUDIT-C, which includes only the first 3 of the 10 AUDIT items. We also obtained cumulative AUD diagnoses, which are also more informative than assessments obtained at a single time point. Because the diagnosis of AUD is based on features other than alcohol consumption *per* se^2,5^, access to the AUD diagnosis from the EHR augmented the information provided by the AUDIT-C phenotype. Although EHR diagnostic data are heterogeneous, large-scale biobanks such as the MVP yield greater statistical power to link genes to health-related traits documented in the EHR than can ordinarily be achieved in prospective studies^23^, justifying the lower resolution of EHR data. However, because the MVP sample is predominantly comprised of EA males, statistical power was limited in both the GWAS and the post-GWAS analyses of other populations and some female samples. Future studies with larger sample sizes are needed to identify additional variation contributing to these alcohol-related traits and to elucidate their interrelationship.

The SNP heritability of our GWASs was lower than that seen in the meta-analysis of the UKBB and 23andMe data^8^. For the AUDIT-C, the estimated SNP heritability was 0.068 in EAs (0.068 in males and 0.099 in females) and 0.062 in AAs. For AUD, the estimated SNP heritability was 0.056 in EAs (0.054 in males and 0.110 in females) and 0.100 in AAs. These estimates may reflect the lower number of SNPs tested in our sample compared with the meta-analysis of UKBB and 23andMe data. The nominally higher SNP heritability in females than males may be due to the substantially smaller size of the female subsample. Although we found no significant difference in PRS between males and females, because of substantially smaller number of women in MVP, there is much less power for the PRS in this subgroup and for comparing the PRS by sex.

Despite these limitations, the large, diverse, and similarly-ascertained sample enabled us to identify multiple novel GWS findings for both AUDIT-C score and AUD diagnosis, and thereby to help elucidate the relationship between drinking level and AUD risk. The large sample provided high power for PRS analyses in other samples, as demonstrated here in the Penn Medicine Biobank and Yale-Penn samples. The genetic differences between the two alcohol-related traits and the observed opposite genetic correlations between them point to potentially important differences in comorbidity and prognosis. Our findings underscore the need to identify the functional effects of the risk variants, especially where they diverge by trait, to elucidate the nature of the trait-related differences. Focusing on variants linked to AUD, but not AUDIT-C, could identify targets for the development of medications to treat the disorder, while variation in AUDIT-C could help in developing interventions to reduce drinking and thereby prevent the morbidity associated with it. The findings reported here could also help to identify individuals at high risk of AUD through the use of PRS. This effort could be augmented using knowledge of the full set of phenotypes that associated with AUD through the use of genetic correlations and PheWASs.

## Methods

The MVP is an observational cohort study and biobank supported by the U.S. Department of Veterans Affairs (VA). Phenotypic data were collected from MVP participants using questionnaires and the VA EHR and a blood sample was obtained for genetic analysis.

**Phenotypes:** AUDIT-C scores and AUD diagnostic codes were obtained from the VA EHR.

<u>The AUDIT-C</u> comprises the first three items of the AUDIT and measures typical quantity (item 1) and frequency (item 2) of drinking and frequency of heavy or binge drinking (item 3). The AUDIT-C is a mandatory annual assessment for all veterans seen in primary care. Our analyses used AUDIT-C data collected from October 1, 2007–February 23, 2017. We validated the phenotype in a sample of 1,851 participants from the Veterans Aging Cohort Study^33^, in which we found a highly significant association of AUDIT-C scores with the plasma concentration of phosphatidylethanol, a direct, quantitative biomarker that is correlated with the level of alcohol consumption. In the AA part of this sample (n=1,503), the AUDIT-C score was highly significantly associated with rs2066702, a missense (Arg369Cys) polymorphism of *ADH1B*, the minor allele of which is common in that population and has been associated with alcohol dependence^25^. We also examined AUDIT-C scores in 167,721 MVP participants (57,677 AAs and 110,044 EAs)^24^, comparing the association of AUDIT-C scores and AUD diagnoses with the frequency of the minor allele of rs2066702 in AAs and rs1229984 (Arg48His) in EAs. Both polymorphisms exert large effects on alcohol metabolism^39^ and are among the genetic variants associated most consistently with alcohol-related traits in both AAs and EAs^8,12,18^. In both populations, we found a stronger association between age-adjusted mean AUDIT-C score and *ADH1B* minor allele frequency than between AUD diagnostic codes and the frequency of the minor alleles^24^. However, because AUD diagnoses accounted for unique variance in the frequency of the minor alleles in both populations, we concluded that the two phenotypes, although correlated, are distinct traits. Thus, in the present study, we used GWAS to examine these traits separately and to adjust for the effects of AUD in the AUDIT-C GWAS and the effects of AUDIT-C in the GWAS of AUD.

As described previously^24^, we calculated the age-adjusted mean AUDIT-C value for each participant using age 50 as the reference point and down-weighting scores for individuals younger than 50 and up-weighting scores for individuals older than 50. There were 272,842 participants in the AUDIT-C analyses. The methods for differentiating the populations are presented below.

### Alcohol Use Disorder (AUD)

The principal classes of alcohol-related disorders in the ICD are alcohol abuse and alcohol dependence. We used ICD-9 codes 303.X (dependence) and 305-305.03 (abuse) and ICD-10 codes F10.1 (abuse) and F10.2 (dependence) to identify subjects diagnosed with either of these disorders, as suggested previously^40^ (see Supplementary Table 51). Participants with at least one inpatient or two outpatient alcohol-related ICD-9/10 codes (N = 274,391) from 2000-2018 were assigned a diagnosis of AUD, an approach that has been shown to yield greater specificity of ICD codes than chart review^41^.

**Genotyping:** MVP GWAS genotyping was performed using an Affymetrix Axiom Biobank Array with 686,693 markers. Subjects or SNPs with genotype call rate < 0.9 or high heterozygosity were removed, leaving 353,948 subjects and 657,459 SNPs for imputation^22^.

### Imputation and Principal Components Analysis (PCA)

Imputation was performed with EAGLE2^42^ to pre-phase each chromosome and Minimac3^43^ to impute genotypes with 1000 Genomes Project phase 3 data^44^ as the reference panel. Subjects with no demographic information or whose genotypic and phenotypic sex did not match were removed. We also removed one subject randomly from each pair of related individuals (kinship coefficient threshold = 0.0884). A greedy algorithm was implemented for network-like relationships among 3 or more individuals, leaving 331,736 subjects for subsequent analyses.

To differentiate population groups, we performed PCA analyses using common SNPs (MAF > 0.05) shared in MVP [pruned using linkage disequilibrium (LD) of *r*^2^ > 0.2] and the 1000 Genomes phase 3 reference panels for European (EUR), African (AFR), admixed American (AMR), East Asian (EAS), and South Asian (SAS) populations using FastPCA in EIGENSOFT^45^. We analyzed 80,871 SNPs in MVP and 1000 Genomes for use in the PCA analyses. The Euclidean distances between each participant and the centers of the 5 reference populations (i.e., across all subjects) were calculated using the first 10 PCs, with each participant assigned to the nearest reference population. A total of 242,317 EA; 61,762 AA; 15,864 Hispanic and Latino American (LA); 1,565 East Asian American (EAA); and 228 South Asian American (SAA) subjects were identified. A second PCA (within each group) yielded the first 10 PCs for each. Participants with PC scores > 3 standard deviations from the mean of any of the 10 PCs were removed as outliers, leaving 209,020 EA; 57,340 AA; 14,425 LA; 1,410 EAA; and 196 SAA subjects. Within genetically defined populations, we calculated population-specific imputation INFO scores using SNPTEST v2^46^ and retained SNPs with INFO scores > 0.7 for association analyses. Imputed genotypes with posterior probability ≥ 0.9 were transferred to best guess.

We removed both genotyped and imputed SNPs with genotype call rates or best guess rates ≤ 0.95 and HWE p-value ≤ 1 × 10^−6^ in each population, using different MAF thresholds to filter SNPs: EA (0.0005), AA (0.001), LA (0.01), EAA (0.05), and SAA (0.05). The approximate number of SNPs remaining in each population was: EA: 6.8 million, AA: 12.5 million, LA: 5.6 million, EAA: 2.6 million, and SA: 2.6 million.

### Genome-wide Association Analyses

Individuals < 22 or > 90 years old and those with missing AUDIT-C scores were removed, leaving 200,680 EAs; 56,495 AAs; 14,112 LAs; 1,366 EAAs; and 189 SAAs in the AUDIT-C GWAS and 202,004 EAs (34,658 cases; 167,346 controls); 56,648 AAs (17,267 cases; 39,381 controls); 14,175 LAs (3,449 cases; 10,726 controls); 1,374 EAAs (164 cases; 1,210 controls); and 190 SAs (44 cases; 144 controls) in the AUD GWAS. We used linear regression for the GWAS of age-adjusted mean AUDIT-C score and logistic regression for AUD diagnosis; in both cases age, sex, and the first 10 PCs were covariates. To evaluate the impact on AUD findings of controlling for AUDIT-C, and the impact on AUDIT-C findings of controlling for AUD, we repeated the GWAS for AUD with AUDIT-C as a covariate and AUDIT-C with AUD as a covariate. For both phenotypes, following GWAS in each of the 5 populations, the summary statistics were combined within phenotype in trans-population meta-analyses. SNPs in EAs or those present in at least two populations were meta-analyzed. Sex-stratified GWAS for both phenotypes were then performed in groups large enough to permit it-EA, AA, LA, and EAA men and EA, AA, and LA women-and the data were meta-analyzed within sex and phenotype. All meta-analyses were performed using a sample-size-weighted scheme that was implemented in METAL^47^.

To identify independent signals in each population, we performed LD clumping using PLINK v1.90b4.4^48^. We identified an index SNP (p < 5 × 10^−8^) with the smallest p-value in a 500-kb genomic window and *r*^2^ < 0.1 with other index SNPs. Because in EAAs there is extended linkage disequilibrium at the *ALDH2* locus, we used a 2,500-kb window in that population. In the chr4q23-q24 region, where we identified multiple apparently independent signals for both AUDIT-C and AUD, we used conditional associations to differentiate independent signals from partially overlapping ones.

<u>Gene-based Association Analysis</u> was performed using Multi-marker Analysis of GenoMic Annotation (MAGMA)^49^, which uses a multiple regression approach to detect multi-marker effects that account for SNP p-values and LD between markers. We used the default setting (no window around genes) to consider 18,575 autosomal genes for the analysis, with p < 2.69 × 10^−6^ (0.05/18,575) considered GWS. For each population, we used the respective population from the 1000 Genomes Project phase 3 as the LD reference.

<u>Pathway and Biological Enrichment Analyses</u> were performed for each population using the FUMA platform^26^, with independent significant SNPs identified using the default settings. Positional gene mapping identified genes up to 10 kb from each independent significant SNP. Hypergeometric tests were used to examine the enrichment of prioritized chemical and genetic perturbation gene sets, canonical pathways, and GO biological processes (obtained from MsigDB c2), and GWAS-catalog enrichment (obtained from reported genes from the GWAS-catalog). We report all significantly enriched gene sets based on a false discovery rate (FDR) adjusted p-value < 0.05.

### Linkage Disequilibrium Score Regression (LDSC)^27^

Population-specific LD scores were calculated based on 1000 Genomes phase 3 datasets according to the LDSC tutorial, using SNPs selected from HapMap 3^50^ after excluding the major histocompatibility complex (MHC) region (chr6:26Mb-34Mb); only ancestry groups with large sample size (N > 10,000) were analyzed using LDSC. Of note, LDSC could be biased in admixed populations because reference panels are not provided for AAs and LAs in that application^27^. We calculated LD scores for 1,215,001 SNPs in EAs; 1,322,841 SNPs in AAs; and 1,243,726 SNPs in LAs. The LDSC analyses used SNPs with imputation INFO ≥ 0.9 in each population and that were LD scored in 1000 Genomes. LD score regression intercepts for available datasets were estimated to distinguish polygenic heritability from inflation. SNP-based heritability 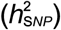 was estimated from GWAS summary statistics for both AUDIT-C and AUD. The sex-specific 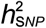 was also estimated in EA males and females, AA males, and LA males.

We estimated partitioned 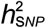 using genomic features or functional categories^28^ for both AUDIT-C and AUD in the largest dataset, EAs, and then tested for enrichment of the partitioned 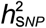 in different annotations. First, we used a baseline model consisting of 53 functional categories, including UCSC gene models [exons, introns, promotors, untranslated regions (UTRs)], ENCODE functional annotations^51^, Roadmap epigenomic annotations^52^, and FANTOM5 enhancers^53^. We then analyzed cell type-specific annotations and identified enrichments of 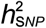 in 10 cell types, including adrenal and pancreas, central nervous system (CNS), cardiovascular, connective tissue and bone, gastrointestinal, immune and hematopoietic, kidney, liver, skeletal muscle and other. Gene expression and chromatin data were also analyzed to identify disease-relevant tissues, cell types, and tissue-specific epigenetic annotations. We used LDSC to test for enriched heritability in regions surrounding genes with the highest tissue-specific expression or with epigenetic marks^29^. Sources of data that were analyzed included 53 human tissue or cell type RNA-seq data from the Genotype-Tissue Expression Project (GTEx)^54^; 152 human, mouse, or rat tissue or cell type array data from the Franke lab^55^; 3 sets of mouse brain cell type array data from Cahoy et al^56^; 292 mouse immune cell type array data from ImmGen^57^; and 396 human epigenetic annotations (6 features in 88 primary cell types or tissues) from the Roadmap Epigenomics Consortium^52^. In the analysis of each trait in each dataset, we used FDR < 0.05 to indicate significant enrichment for the 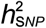.

### Genetic Correlations (*r_g_*)

We estimated genetic correlations between AUDIT-C and AUD (from MVP), and with other traits in LD Hub^58^ or from published studies using LDSC, which is robust to sample overlap^30^. First, we estimated the *r_g_* between AUDIT-C and AUD using the summary data generated in this study. AAs, EAs and LAs were analyzed separately using the corresponding 1000 Genome phase3 population as reference. Genetic correlations in EAA and SAA were not analyzed. The *r*_g_ between AUDIT-C and AUD was out of bounds due to the 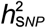 not differing significantly from zero. Then we tested the *r_g_* between males and females within each trait. We estimated the *r_g_* for AUDIT-C and AUD with 216 published traits in LD Hub and 493 unpublished traits from the UK Biobank. We resolved redundancy in phenotypes by manually selecting the published version of the phenotype or using the largest sample size. We also calculated the genetic correlations of both AUDIT-C and AUD with 5 traits for which GWAS were recently published or posted, including anorexia nervosa^59^, alcohol dependence^13^, attention deficit hyperactivity disorder^60^, autism spectrum disorder^61^ and major depressive disorder (summary data without the 23andMe sample)^62^, bringing the total number of tested traits to 714. A Bonferroni correction was applied separately for AUDIT-C and AUD, and traits with a corrected p-value < 0.05 were considered significantly correlated. Because the results were similar whether the intercept was constrained or not, we present here the original results without constraint.

### Polygenic Risk Scores (PRS) in the MVP

To generate PRS from GWAS summary statistics in the MVP sample, we first conducted a GWAS for AUDIT-C and AUD as above, but restricted our analysis to two thirds of the total sample by splitting the total sample randomly, keeping the number of AUD cases/controls balanced in each part (EA: N=139,346, AA: N=38,226). GWS loci identified from this analysis were the same as those in the larger sample, but slightly decreased in significance. PRS were generated for the remaining EAs (N=69,674) and AAs (N=19,114)_as the sum of all variants carried, weighted by the effect size of the variant in the GWAS. PRS were generated using PLINK2^63^. We performed p-value informed clumping with a distance threshold of 250 kb and *r*^2^ = 0.1. Risk scores were calculated for a range of p-value thresholds (*P* ≤ 1 × 10^−7^, 1 × 10^−6^, 1 × 10^−5^, 1 × 10^−4^, 1 × 10^−3^, 0.01, 0.05, 0.5, 1.0). PRS were standardized with mean = 0 and SD = 1. Logistic regression was used to test for association with AUDIT-C and AUD phenotypes, with PRS as the independent variable and AUDIT-C or AUD as the dependent variable, with age, sex, and the first 5 PCs as covariates.

### PRS in the Penn Medicine BioBank (PMBB)

Population-specific summary statistics from the AUDIT-C and AUD GWAS in MVP were used to generate PRS in the PMBB, an independent sample. PRS were generated for EAs (N = 8,524) and AAs (N = 2,031) as above using the PRSice2 package^64^ with imputed allele dosage as the target dataset. As recommended in the software, we performed p-value informed clumping with a distance threshold of 250 kb and an *r*^2^ = 0.1. We excluded the MHC region. Risk scores were calculated for a range of p-value thresholds (*P* ≤ 1 × 10^−7^, 1 × 10^−6^, 1 × 10^−5^, 1 × 10^−4^, 1 × 10^−3^, 0.01, 0.05, 0.5, 1.0) and standardized with mean = 0 and SD = 1. To identify individuals with alcohol-related disorders, we utilized phecodes, a method to aggregate ICD codes^65^. First, we extracted ICD-9 and ICD-10 data for 48,610 individuals from the EHR. To facilitate mapping to phecodes, ICD-10 codes were back converted to ICD-9 using 2017 general equivalency mapping (GEM). The ICD-10 conversions were combined with the ICD-9 codes to create a dataset with 10,682 unique ICD-9 codes. ICD-9 codes were aggregated to phecodes using the PheWAS R package^65^ to create 1,812 phecodes. Individuals are considered cases for the phenotype if they had at least 2 instances of the phecode, controls if they had no instance of the phecode, and “other/missing” if they had one instance or a related phecode. Logistic regression was used to test the association of the PRS with the alcohol-related disorders phecode (phecode number 317) and its sub-phenotype, alcoholism (phecode number 317.1). The analysis was performed in R with PRS as the independent variable and diagnosis as the dependent variable and age, sex, and the first 10 PCs as covariates.

### PRS in the Yale-Penn cohort

We also tested the PRS of AUDIT-C and AUD for DSM-IV alcohol dependence criterion counts in the Yale-Penn cohort^25^. There are three phases of the Yale-Penn sample: phase 1 contains 3,110 AAs and 1,718 EAs exposed to alcohol; phase 2 contains 1,667 AAs and 1,689 EAs exposed to alcohol; phase3 contains 556 AAs and 999 EAs exposed to alcohol. PRS were generated for EAs and AAs in each phase as described above and risk scores were calculated for a range of p-value thresholds (p ≤ 1 × 10^−7^, 1 × 10^−6^, 1 × 10^−5^, 1 × 10^−4^, 1 × 10^−3^, 0.01, 0.05, 0.5, 1.0). Different from PRS in MVP and PMBB, to correct for the relatedness in the Yale-Penn subjects, a linear mixed model implemented in GEMMA^66^ was used to test the association between PRS score and DSM-IV alcohol dependence criterion counts, with age, sex, and the first 10 PCs as covariates. Meta-analyses of data from the three phases were performed in AAs (N = 5,333) and EAs (N = 4,406) separately.

### Phenome-wide Association Analysis (PheWAS)

We extracted ICD-9 data from the EHR for 353,323 genotyped veterans. Of these, 277,531 individuals had 2 or more separate encounters in the VA Healthcare System in each of the two years prior to enrollment in MVP, consisting of 21,209,658 records. ICD-9 codes were aggregated to phecodes using the PheWAS R package to create 1,812 phecodes. To improve the specificity of these codes, individuals with at least 2 instances of the phecode were considered cases, those with no instance of the phecode controls, and those with one instance of a phecode or a related phecode as “other.” A PheWAS using logistic regression models with either AUDIT-C or AUD PRS as the independent variable, phecodes as the dependent variables, and age, sex and the first 5 PCs as covariates, were used to identify secondary phenotypic associations. A phenome-wide significance threshold of 2.96 × 10^−5^ was applied to account for multiple testing.

### GWAS Adjusted for BMI

As described below, for both alcohol-related traits, we identified a GWS SNP in *FTO*, variation in which has been associated with BMI and risk of obesity^67^. To examine whether BMI confounded the association with this and other loci and the genetic correlations with other traits, we repeated the GWAS for AUDIT-C and AUD using BMI as an additional covariate. Data on BMI were from the MVP baseline survey and the EHR. For AUDIT-C, 200,092 EAs; 56,239 AAs; 14,029 LAs; 1,352 EAAs; and 185 SAAs had BMI data available. For AUD, 201,320 EAs; 56,347 AAs; 14,075 LAs; 1,360 EAAs; and 186 SAAs had BMI data available. After GWAS, we analyzed the genetic correlations between BMI-adjusted traits and other publicly available traits (N = 714), with Bonferroni correction for multiple testing.

## Supporting information

link to supplemental figures

## References

1 World Health Organization. Global Status Report on Alcohol and Health. Geneva: WHO. http://www.who.int/substance_abuse/publications/global_alcohol_report/en/; accessed June 5, 2018. (2014).

2 American Psychiatric Association. Diagnostic and statistical manual of mental disorders (5th ed.) Arlington, VA: American Psychiatric Association. (2013).

3 American Psychiatric Association. Diagnostic and statistical manual of mental disorders: DSM-IV-TR. Washington, DC: American Psychiatric Association. (2000).

4 Saunders, J.B., Aasland, O.G., Babor, T.F., de la Fuente, J.R. & Grant, M. Development of the Alcohol Use Disorders Identification Test (AUDIT): WHO Collaborative Project on Early Detection of Persons with Harmful Alcohol Consumption–II. Addiction 88, 791–804 (1993).

5 Babor, T.F. et al. AUDIT: The Alcohol Use Disorders Identification Test : guidelines for use in primary health care. 2nd ed. Geneva : World Health Organization. (2001).

6 Mbarek, H. et al. The genetics of alcohol dependence: Twin and SNP-based heritability, and genome-wide association study based on AUDIT scores. Am J Med Genet B Neuropsychiatr Genet 168, 739–748, doi:10.1002/ajmg.b.32379 (2015).

7 Sanchez-Roige, S. et al. Genome-wide association study of alcohol use disorder identification test (AUDIT) scores in 20 328 research participants of European ancestry. Addict Biol, doi:10.1111/adb.12574 (2017).

8 Sanchez-Roige, S. et al. Genome-wide association study meta-analysis of the Alcohol Use Disorders Identification Test (AUDIT) in two population-based cohorts. Am J Psychiatry, appiajp201818040369, doi:10.1176/appi.ajp.2018.18040369 (2018).

9 Verhulst, B., Neale, M.C. & Kendler, K.S. The heritability of alcohol use disorders: a meta-analysis of twin and adoption studies. Psychol Med 45, 1061–1072, doi:10.1017/S0033291714002165 (2015).

10 Vrieze, S.I., McGue, M., Miller, M.B., Hicks, B.M. & Iacono, W.G. Three mutually informative ways to understand the genetic relationships among behavioral disinhibition, alcohol use, drug use, nicotine use/dependence, and their co-occurrence: twin biometry, GCTA, and genome-wide scoring. Behav Genet 43, 97–107, doi:10.1007/s10519-013-9584-z (2013).

11 Yang, C. et al. Exploring the genetic architecture of alcohol dependence in African-Americans via analysis of a genomewide set of common variants. Hum Genet 133, 617–624, doi:10.1007/s00439-013-1399-8 (2014).

12 Hart, A.B. & Kranzler, H.R. Alcohol dependence genetics: Lessons learned from genome-wide association studies (GWAS) and post-GWAS analyses. Alcohol Clin Exp Res 39, 1312–1327, doi:10.1111/acer.12792 (2015).

13 Walters, R.K. et al. Trans-ancestral GWAS of alcohol dependence reveals common genetic underpinnings with psychiatric disorders. bioRxiv. doi:https://doi.org/10.1101/257311 (2018).

14 Schumann, G. et al. Genome-wide association and genetic functional studies identify autism susceptibility candidate 2 gene (AUTS2) in the regulation of alcohol consumption. Proc Natl Acad Sci U S A 108, 7119–7124, doi:10.1073/pnas.1017288108 (2011).

15 Takeuchi, F. et al. Confirmation of ALDH2 as a major locus of drinking behavior and of its variants regulating multiple metabolic phenotypes in a Japanese population. Circ J 75, 911–918 (2011).

16 Kapoor, M. et al. A meta-analysis of two genome-wide association studies to identify novel loci for maximum number of alcoholic drinks. Hum Genet 132, 1141–1151, doi:10.1007/s00439-013-1318-z (2013).

17 Quillen, E.E. et al. ALDH2 is associated to alcohol dependence and is the major genetic determinant of “daily maximum drinks” in a GWAS study of an isolated rural Chinese sample. Am J Med Genet B Neuropsychiatr Genet 165B, 103–110, doi:10.1002/ajmg.b.32213 (2014).

18 Xu, K. et al. Genomewide association study for maximum number of alcoholic drinks in European Americans and African Americans. Alcohol Clin Exp Res 39, 1137–1147, doi:10.1111/acer.12751 (2015).

19 Gelernter, J. et al. Genomewide association study of alcohol dependence and related traits in a Thai population. Alcohol Clin Exp Res 42, 861–868, doi:10.1111/acer.13614 (2018).

20 Schumann, G. et al. KLB is associated with alcohol drinking, and its gene product beta-Klotho is necessary for FGF21 regulation of alcohol preference. Proc Natl Acad Sci U S A 113, 14372–14377, doi:10.1073/pnas.1611243113 (2016).

21 Clarke, T.K. et al. Genome-wide association study of alcohol consumption and genetic overlap with other health-related traits in UK Biobank (N=112 117). Mol Psychiatry 22, 1376–1384, doi:10.1038/mp.2017.153 (2017).

22 Gaziano, J.M. et al. Million Veteran Program: A mega-biobank to study genetic influences on health and disease. J Clin Epidemiol 70, 214–223, doi:10.1016/j.jclinepi.2015.09.016 (2016).

23 Collins, R. What makes UK Biobank special? Lancet 379, 1173–1174, doi:10.1016/S0140-6736(12)60404-8 (2012).

24 Justice, A.C. et al. AUDIT-C and ICD codes as phenotypes for harmful alcohol use: association with ADH1B polymorphisms in two US populations. Addiction 113, 2214–2224, doi:10.1111/add.14374 (2018).

25 Gelernter, J. et al. Genome-wide association study of alcohol dependence:significant findings in African- and European-Americans including novel risk loci. Mol Psychiatry 19, 41–49, doi:10.1038/mp.2013.145 (2014).

26 Watanabe, K., Taskesen, E., van Bochoven, A. & Posthuma, D. Functional mapping and annotation of genetic associations with FUMA. Nat Commun 8, 1826, doi:10.1038/s41467-017-01261-5 (2017).

27 Bulik-Sullivan, B.K. et al. LD Score regression distinguishes confounding from polygenicity in genome-wide association studies. Nat Genet 47, 291–295, doi:10.1038/ng.3211 (2015).

28 Finucane, H.K. et al. Partitioning heritability by functional annotation using genome-wide association summary statistics. Nat Genet 47, 1228–1235, doi:10.1038/ng.3404 (2015).

29 Finucane, H.K. et al. Heritability enrichment of specifically expressed genes identifies disease-relevant tissues and cell types. Nat Genet 50, 621–629, doi:10.1038/s41588-018-0081-4 (2018).

30 Bulik-Sullivan, B. et al. An atlas of genetic correlations across human diseases and traits. Nat Genet 47, 1236–1241, doi:10.1038/ng.3406 (2015).

31 Grant, J. D. et al. Alcohol consumption indices of genetic risk for alcohol dependence. Biol Psychiatry 66, 795–800, doi:10.1016/j.biopsych.2009.05.018 (2009).

32 Kendler, K.S., Myers, J., Dick, D. & Prescott, C.A. The relationship between genetic influences on alcohol dependence and on patterns of alcohol consumption. Alcohol Clin Exp Res 34, 1058–1065, doi:10.1111/j.1530-0277.2010.01181.x (2010).

33 Justice, A.C. et al. Validating harmful alcohol use as a phenotype for genetic discovery using phosphatidylethanol and a polymorphism in ADH1B. Alcohol Clin Exp Res 41, 998–1003, doi:10.1111/acer.13373 (2017).

34 Wood, A.M. et al. Risk thresholds for alcohol consumption: combined analysis of individual-participant data for 599 912 current drinkers in 83 prospective studies. Lancet 391, 1513–1523, doi:10.1016/S0140-6736(18)30134-X (2018).

35 Chou, S.P. et al. Alcohol use disorders, nicotine dependence, and co-occurring mood and anxiety disorders in the United States and South Korea-a cross-national comparison. Alcohol Clin Exp Res 36, 654–662, doi:10.1111/j.1530-0277.2011.01639.x (2012).

36 Lai, H.M., Cleary, M., Sitharthan, T. & Hunt, G.E. Prevalence of comorbid substance use, anxiety and mood disorders in epidemiological surveys, 1990-2014: A systematic review and meta-analysis. Drug Alcohol Depend 154, 1–13, doi:10.1016/j.drugalcdep.2015.05.031 (2015).

37 Kendler, K.S., Ohlsson, H., Sundquist, J. & Sundquist, K. School achievement, IQ, and risk of alcohol use disorder: A prospective, co-relative analysis in a Swedish national cohort. J Stud Alcohol Drugs 78, 186–194 (2017).

38 Eyawo, O. et al. Alcohol and mortality: Combining self-reported (AUDIT-C) and biomarker detected (PEth) alcohol measures among HIV infected and uninfected. J Acquir Immune Defic Syndr 77, 135–143, doi:10.1097/QAI.0000000000001588 (2018).

39 Polimanti, R. & Gelernter, J. ADH1B: From alcoholism, natural selection, and cancer to the human phenome. Am J Med Genet B Neuropsychiatr Genet 177, 113–125, doi:10.1002/ajmg.b.32523 (2018).

40 Piette, J.D., Barnett, P.G. & Moos, R.H. First-time admissions with alcohol-related medical problems: a 10-year follow-up of a national sample of alcoholic patients. J Stud Alcohol 59, 89–96 (1998).

41 Justice, A.C. et al. Medical disease and alcohol use among veterans with human immunodeficiency infection: A comparison of disease measurement strategies. Med Care 44, S52–60, doi:10.1097/01.mlr.0000228003.08925.8c (2006).

42 Loh, P.R. et al. Reference-based phasing using the Haplotype Reference Consortium panel. Nat Genet 48, 1443–1448, doi:10.1038/ng.3679 (2016).

43 Browning, B.L. & Browning, S.R. Genotype imputation with millions of reference samples. Am J Hum Genet 98, 116–126, doi:10.1016/j.ajhg.2015.11.020 (2016).

44 1000 Genomes Project Consortium et al. A global reference for human genetic variation. Nature 526, 68–74, doi:10.1038/nature15393 (2015).

45 Galinsky, K.J. et al. Fast principal-component analysis reveals convergent evolution of ADH1B in Europe and East Asia. Am J Hum Genet 98, 456–472, doi:10.1016/j.ajhg.2015.12.022 (2016).

46 Marchini, J. & Howie, B. Genotype imputation for genome-wide association studies. Nat Rev Genet 11, 499–511, doi:10.1038/nrg2796 (2010).

47 Willer, C.J., Li, Y. & Abecasis, G.R. METAL: fast and efficient meta-analysis of genomewide association scans. Bioinformatics 26, 2190–2191, doi:10.1093/bioinformatics/btq340 (2010).

48 Purcell, S. et al. PLINK: a tool set for whole-genome association and population-based linkage analyses. Am J Hum Genet 81, 559–575, doi:10.1086/519795 (2007).

49 de Leeuw, C.A., Mooij, J.M., Heskes, T. & Posthuma, D. MAGMA: generalized gene-set analysis of GWAS data. PLoS Comput Biol 11, e1004219, doi:10.1371/journal.pcbi.1004219 (2015).

50 International HapMap Consortium et al. Integrating common and rare genetic variation in diverse human populations. Nature 467, 52–58, doi:10.1038/nature09298 (2010).

51 Encode Project Consortium. An integrated encyclopedia of DNA elements in the human genome. Nature 489, 57–74, doi:10.1038/nature11247 (2012).

52 Roadmap Epigenomics Consortium et al. Integrative analysis of 111 reference human epigenomes. Nature 518, 317–330, doi:10.1038/nature14248 (2015).

53 Andersson, R. et al. An atlas of active enhancers across human cell types and tissues. Nature 507, 455–461, doi:10.1038/nature12787 (2014).

54 GTEx Consortium. Human genomics. The Genotype-Tissue Expression (GTEx) pilot analysis: multitissue gene regulation in humans. Science 348, 648–660, doi:10.1126/science.1262110 (2015).

55 Fehrmann, R.S. et al. Gene expression analysis identifies global gene dosage sensitivity in cancer. Nat Genet 47, 115–125, doi:10.1038/ng.3173 (2015).

56 Cahoy, J.D. et al. A transcriptome database for astrocytes, neurons, and oligodendrocytes: a new resource for understanding brain development and function. J Neurosci 28, 264–278, doi:10.1523/JNEUROSCI.4178-07.2008 (2008).

57 Heng, T.S., Painter, M.W. & Immunological Genome Project, C. The Immunological Genome Project: networks of gene expression in immune cells. Nat Immunol 9, 1091–1094, doi:10.1038/ni1008-1091 (2008).

58 Zheng, J. et al. LD Hub: a centralized database and web interface to perform LD score regression that maximizes the potential of summary level GWAS data for SNP heritability and genetic correlation analysis. Bioinformatics 33, 272–279, doi:10.1093/bioinformatics/btw613 (2017).

59 Duncan, L. et al. Significant locus and metabolic genetic correlations revealed in genome-wide association study of anorexia nervosa. Am J Psychiatry 174, 850–858, doi:10.1176/appi.ajp.2017.16121402 (2017).

60 Demontis, D. et al. Discovery of the first genome-wide significant risk loci for ADHD. bioRxiv. doi:https://doi.org/10.1101/145581 (2017).

61 Autism Spectrum Disorders Working Group of The Psychiatric Genomics Consortium. Meta-analysis of GWAS of over 16,000 individuals with autism spectrum disorder highlights a novel locus at 10q24.32 and a significant overlap with schizophrenia. Mol Autism 8, 21, doi:10.1186/s13229-017-0137-9 (2017).

62 Wray, N.R. et al. Genome-wide association analyses identify 44 risk variants and refine the genetic architecture of major depression. Nat Genet 50, 668–681, doi:10.1038/s41588-018-0090-3 (2018).

63 Chang, C.C. et al. Second-generation PLINK: rising to the challenge of larger and richer datasets. Gigascience 4, 7, doi:10.1186/s13742-015-0047-8 (2015).

64 Euesden, J., Lewis, C. M. & O’Reilly, P.F. PRSice: Polygenic Risk Score software. Bioinformatics 31, 1466–1468, doi:10.1093/bioinformatics/btu848 (2015).

65 Carroll, R. J., Bastarache, L. & Denny, J. C. R PheWAS: data analysis and plotting tools for phenome-wide association studies in the R environment. Bioinformatics 30, 2375–2376, doi:10.1093/bioinformatics/btu197 (2014).

66 Zhou, X. & Stephens, M. Efficient multivariate linear mixed model algorithms for genome-wide association studies. Nat Methods 11, 407–409, doi:10.1038/nmeth.2848 (2014).

67 Loos, R. J. & Yeo, G. S. The bigger picture of FTO: the first GWAS-identified obesity gene. Nat Rev Endocrinol 10, 51–61, doi:10.1038/nrendo.2013.227 (2014).

